# gmxapi: a high-level interface for advanced control and extension of molecular dynamics simulations

**DOI:** 10.1101/306043

**Authors:** M. Eric Irrgang, Jennifer M. Hays, Peter M. Kasson

## Abstract

**Summary:** Molecular dynamics simulations have found use in a wide variety of biomolecular applications, from protein folding kinetics to computational drug design to refinement of molecular structures. Two areas where users and developers frequently need to extend the built-in capabilities of most software packages are implementing custom interactions, for instance biases derived from experimental data, and running ensembles of simulations. We present a Python high-level interface for the popular simulation package GROMACS that 1) allows custom potential functions without modifying the simulation package code, 2) maintains the optimized performance of GROMACS, and 3) presents an abstract interface to building and executing computational graphs that allows transparent low-level optimization of data flow and task placement. Minimal dependencies make this integrated API for the GROMACS simulation engine simple, portable, and maintainable. We demonstrate this API for experimentally-driven refinement of protein conformational ensembles.

**Availability:** Source and installation instructions are available at https://github.com/kassonlab/gmxapi.

## 1 Introduction

As biomolecular simulations have advanced in complexity and scale, programmatic control of simulations has become a common mode of use. This has been accomplished both through middleware layers (Balasubramanian, et al., 2016; Pronk, et al., 2011) and native programming interfaces (Eastman, et al., 2013; Phillips, et al., 2005), with Python interfaces becoming increasingly common due to Python’s popularity in the scientific computing community, its robust scripting interface, and the rich ecosystem of data analysis and visualization tools available. Among major molecular dynamics (MD) software packages, the few that offer native Python interfaces tend to do so via procedural calls so that the resulting code is executed in a linear, stepwise fashion. This is a natural programming paradigm for users accustomed to writing shell scripts, but it prevents more advanced task placement and parallelization strategies. Packages such as TensorFlow (Abadi, et al., 2016) or the MD overlay software Copernicus (Pronk, et al., 2011) demonstrate an alternative paradigm where the API provides an interface for constructing a *computational task graph* that can then be executed in an optimized manner by the underlying software.

Our design approach is to provide a native interface to the GROMACS MD engine (Pronk, et al., 2013) that supports two common use patterns that require either middleware packages or custom modification of the GROMACS source. This interface also allows simple, intuitive construction of computational task graphs in a manner that permits abstraction of parallel optimizations and ultimately is compatible with advanced machine-learning packages such as TensorFlow to permit mixing of molecular simulation and machine-learning operations.

We therefore present a native Python API to the GROMACS simulation engine that implements these features. Users may drive simulations from Python via simple high-level procedural commands, a more granular object-oriented interface, or through their own extension code. We include a framework for extending GROMACS with MD plugin modules for which Python interfaces are automatically generated, permitting developers to customize GROMACS without modifying the source. The current interface focuses on the MD engine itself; future versions will encompass analysis tools and facilitate integration with third-party analysis software. Here, we outline key features of gmxapi and demonstrate its utility by implementing restrained-ensemble MD simulations for hybrid refinement of protein structures based on experimental data.

## 2 Methods

The gmxapi package consists of a high-level interface in pure Python with a lower-level API implemented as a C++ extension. The Python component provides the gmx module as a stable external interface. Bindings to the libgmxapi C++ API are provided in the submodule gmx.core. C++ implementations for different compute platforms (cloud platforms, GPUs, parallel architectures) may be coded differently but are presented through a consistent interface that abstracts these details away.

**Fig. 1:**
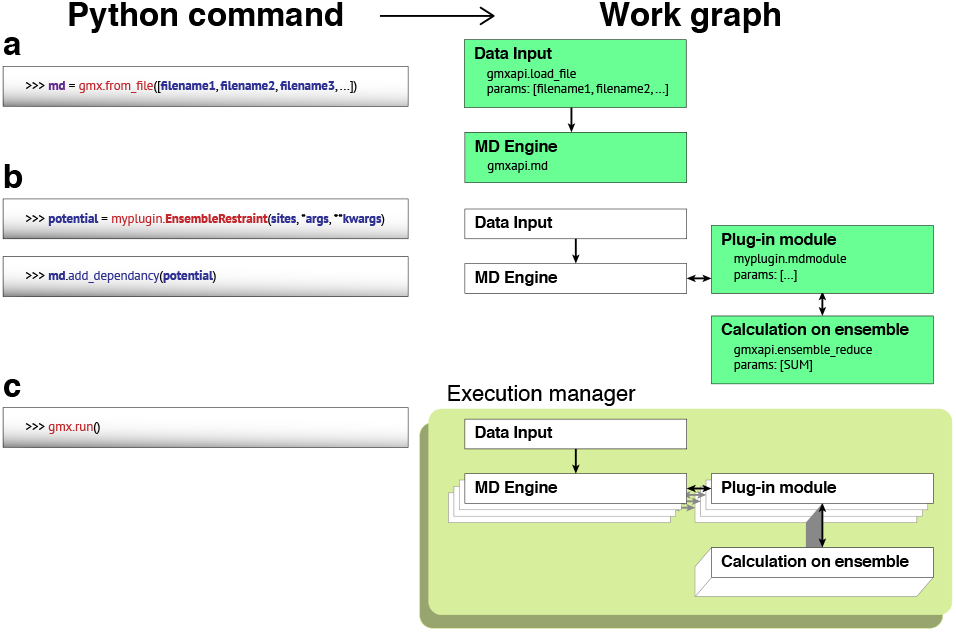
Restrained-ensemble simulations using gmxapi. Schematized are the Python commands to declare an array of MD simulations, bind a custom potential, and run, and the corresponding computational graph.

The high-level Python interface provides essential abstractions for workflow construction and execution. Procedural commands initialize and construct a workflow, which may be serial or parallel at both the individual-simulation and simulation-ensemble levels. Once the workflow is fully described, a single Python function discovers and allocates computing resources and hands off the work specification to an execution manager that translates it into a task graph and executes it.

A plugin API is included to allow custom extensions of GROMACS MD potentials without recompiling the MD engine code itself. Plugins are constructed via C++ templating and Python bindings via pybindll (sample code is provided). Users can thus build a custom plugin and add it to the work specification, and the gmxapi execution manager will bind the custom code into the MD loop at runtime. The result is a friendly Python interface for custom extensions that nonetheless maintains native GROMACS performance. Details on work specification grammar and the plugin interface are given in the Supplementary Data.

## 3 Results

To demonstrate the power of the gmxapi package, we have tested it on restrained-ensemble refinement of protein conformational ensembles using experimental DEER spectroscopy data. This approach, originally published and implemented using CHARMM (Roux and Islam, 2013), is a common workflow in our group using GROMACS that requires custom code in three places: user-specified biasing forces in the core MD engine, analysis code to process predicted ensemble data and update the biasing forces, and parallelization scripts to manage execution, analysis, and data exchange between many ensemble members simultaneously. We have replaced all three of these using the gmxapi plugin interface and simple high-level calls to the gmxapi Python API.

At a high level, restrained-ensemble simulations compute population properties from a set of molecular dynamics simulations, compare those to an experimental measurement, and compute biases to bring the simulated ensemble in better agreement with the experimental one. The experimental data we use are residue-residue distance distributions measured via double electron-electron resonance (DEER) spectroscopy. The simulation algorithm is thus to compute a distance histogram from the estimated ensemble, compare to the measured ensemble, and calculate a distance-dependent biasing force for the simulations, which are run for an interval Δt before the process is repeated (see Supplement). Multiple DEER restraints can be applied in a single simulated ensemble.

The gmxapi calls required to set up a restrained-ensemble workflow are schematized in Fig. 1, and the full source is given in the Supplementary Data. Prior to gmxapi, our group wrote a custom implementation of restrained-ensemble simulations for GROMACS that required 6984 lines of code. With gmxapi our C++ plugin and python source are 127 lines once include and comment statements are excluded. Performance using the external plugin is within 5% of the custom implementation where restraint forces are deeply embedded in the GROMACS code.

The gmxapi package thus provides a high-level interface for the GROMACS MD engine and enables custom plugins for user-specified forces, abstraction of computational context in a task-graph architecture, and first-class management of simulation ensembles. Further improvements will expand this API to cover parallel analysis tasks as well.

## Acknowledgements

We thank Mark Abraham, Michael Shirts, Shantenu Jha, and members of the GROMACS core developer and MolSSI communities for helpful discussions.

## References

Abadi, M., et al. Tensorflow: Large-scale machine learning on heterogeneous distributed systems. arXivpreprint arXiv:l 603.04467 2016.

Balasubramanian, V., et al. Extasy: Scalable and flexible coupling of md simulations and advanced sampling techniques. In, e-Science (e-Science), 2016 IEEE 12th International Conference on. IEEE; 2016. p. 361-370.

Eastman, P., et al. OpenMM 4: A Reusable, Extensible, Hardware Independent Library for High Performance Molecular Simulation. J. Chem. Theory Comput. 2013;9(1):461-469.

Phillips, J.C., et al. Scalable molecular dynamics with NAMD. J Comput Chem 2005;26(16): 1781-1802.

Pronk, S., et al. Copernicus: A new paradigm for parallel adaptive molecular dynamics. Proceedings of 2011 International Conference for High Performance Computing, Networking, Storage and Analysis 2011:60.

Pronk, S., et al. GROMACS 4.5: a high-throughput and highly parallel open source molecular simulation toolkit. Bioinformatics 2013;29(7):845-854.

Roux B. and Islam S.M. Restrained-ensemble molecular dynamics simulations based on distance histograms from double electron-electron resonance spectroscopy. J Phys Chem B 2013;117(17):4733-4739.

